# WITOD: A Tool for Within-Taxon Operational Taxonomic Unit Diversity Analysis

**DOI:** 10.1101/813444

**Authors:** John Kevin Cava, Gaoyang Li, Wei Du, Huansheng Cao

## Abstract

**Motivation:** Taxonomic analysis of microbiomes based on 16S rRNA amplicons has so far been usually restricted to abundance analysis of operational taxonomic units (OTUs), which are usually mapped to genus, as the furthest level for a balance between accuracy and speed. Biodiversity within taxa has been little studied, due to lack of proper computational tools. Within-taxon diversity reflects genetic polymorphism within the taxa to reveal whether there are potentially more variants/lineages for some microbiome-associated traits, e.g., diseases or bioconversion efficiency. The diversity will aid diagnosis decisions.

**Results:** Here we introduce a tool, WITOD, for the WIthin-Taxon Operational taxonomic unit Diversity analysis. WITOD works on the alignment of all the OTU sequences within a taxon to get a non-redundant alignment with consensus regions on both ends (due to the conservation of the 16S regions used). Then the relative abundance of identical OTUs is combined; more specific taxonomy is obtained through BLASTn of these OTU sequences against the Silva database. One of the outputs is an OTU table with these unique OTUs with new taxonomy, combined relative abundance. In addition, another table can be constructed which we call the diversity table that associates the number of OTUs for a specific Taxa and the relative abundance in that environment. This can be used to further discover the importance of differing diversity in a given environment.

**Conclusions:** In this paper, we have introduced a python program WITOD which is a computational tool to analyze the within-taxon biodiversity of microbiome composition. It will be useful in revealing the genetic polymorphism in general and aiding identifying the causative human pathogens or environmental disease vectors. WITOD is available at https://github.com/johncava/WITOD

## Background

In microbiome research, current analysis of 16S rRNA gene amplicons is generally focused on the abundance of operational taxonomical units (OTUs) and the taxa which they belong to (e.g., Guo et al. 2018). However, the biodiversity within taxa has been rarely studied, although it is important to both general microbiome compositional structure dissection and accurate diagnosis of human pathogens (Faith et al. 2015). When genetic polymorphism occurs, these different genotypes are likely to have different phenotypes, which may display different virulence in the case of human pathogens or degrading capabilities among bioengineering species (Ward et al. 2016).One recent study explored the diversity of main water-bloom cyanobacteria and found that abundant genera have high biodiversity (as the number of unique OTUs) (Zhu et al. 2019).Given the importance of the understudied issue, a computational tool needed to be developed.

Technically, this question can be solved by analyzing the clustered OTU sequences generated from popular programs such as QIIME (Caporaso et al 2010) and MOTHUR (Schloss et al 2009). However, the OTU sequences differ in either the length (the number of bases) or the sequence (the order of the bases) of OTU sequences, so some OTU sequences could well be subsequences of longer OTU sequences in the same taxa (e.g., Zhu et al. 2019). Additionally, the 16S rRNA amplicons and thus the resulting OTU sequences are usually short, e.g., about 200 bp (the V3 region) or 460 bp (the V3-V4 region) (Klindworth et al. 2013), these variable regions need to be analyzed with reliable methods for within-taxon diversity (Chakravorty et al. 2007). To reliably solve this issue, OTU sequence alignment should be used, as these OTU sequences are amplified with conserved primers, so the two flanking ends of OTU sequence alignments of the same taxa should have high level of conservation and thus consensus bases. Once these consensus regions are identified, the region between them will be have the same lengths and the number of variant bases in this region is a measure of the within-taxon biodiversity.

## Results

To test our methodology, we take data from a cyanobacterial bloom microbiome at the BUOY site in Harsha Lake (Ohio, USA) (Zhu et al 2019). We then utilize our framework on the OTU table from the BUOY site, and create diversity tables for each sample. From these diversity tables, we are then able to plot how the number of unique OTUs compare to the relative abundance of that OTU in that particular sample. In Figure 4, we plot a 6 different samples from the BUOY site. In addition, each sample has differing equations for each slope.

**Figure 1.**
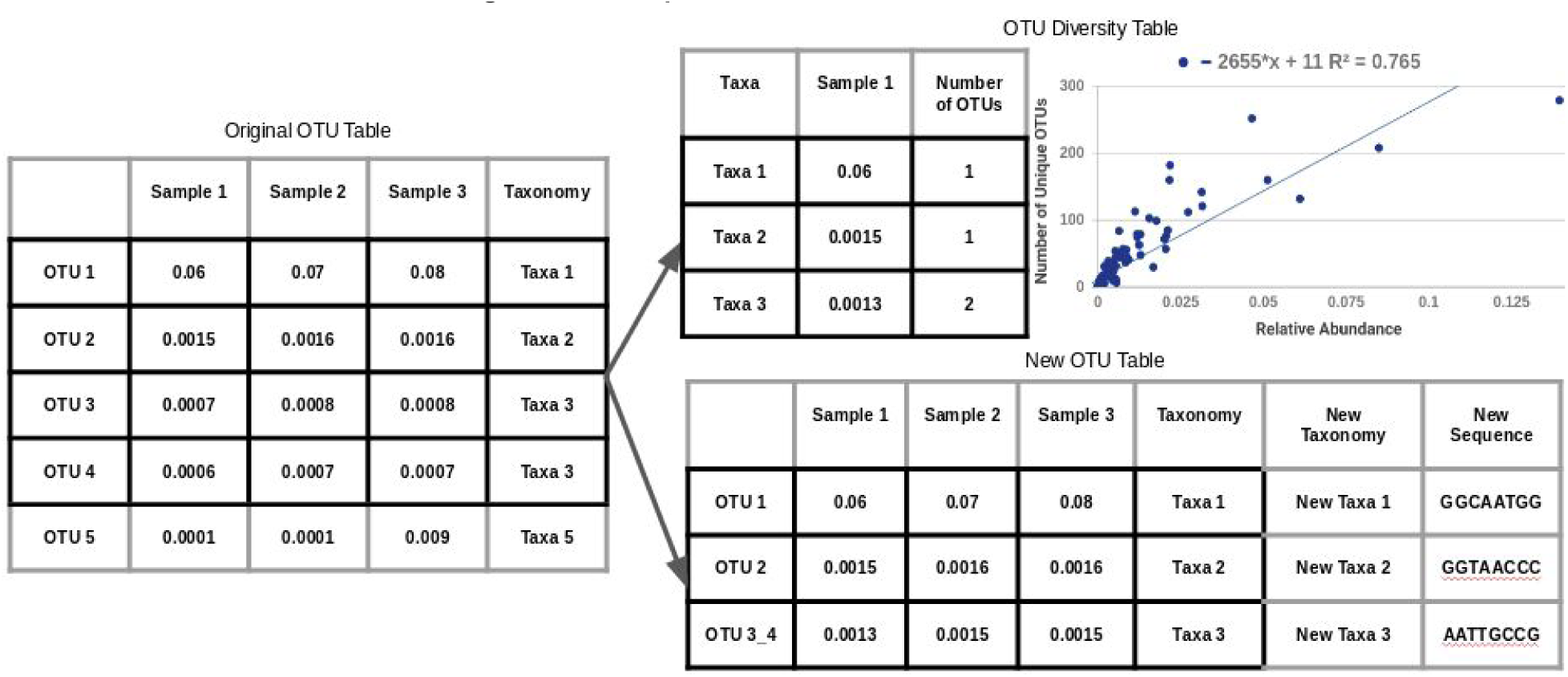
High level description of WITOD.

**Figure 2.**
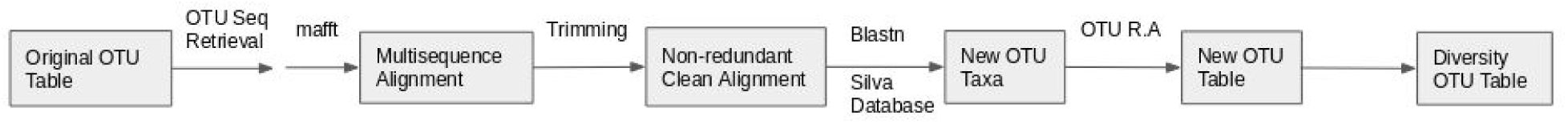
The overall pipeline of WITOD.

**Figure 3.**
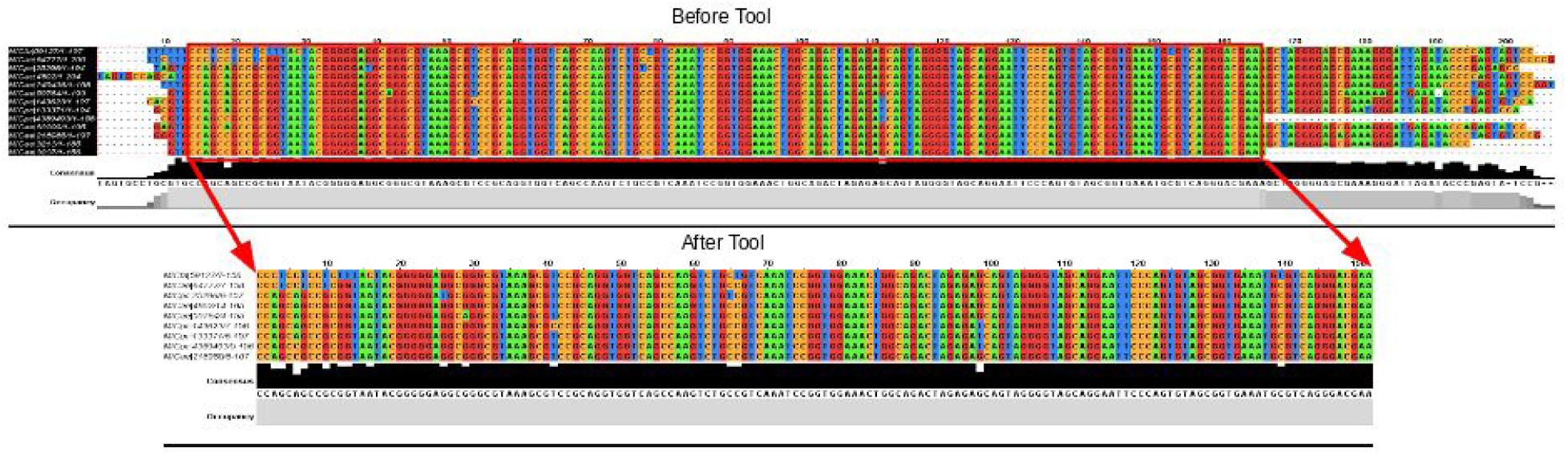
The OTU sequences after trimming.

**Figure 4.**
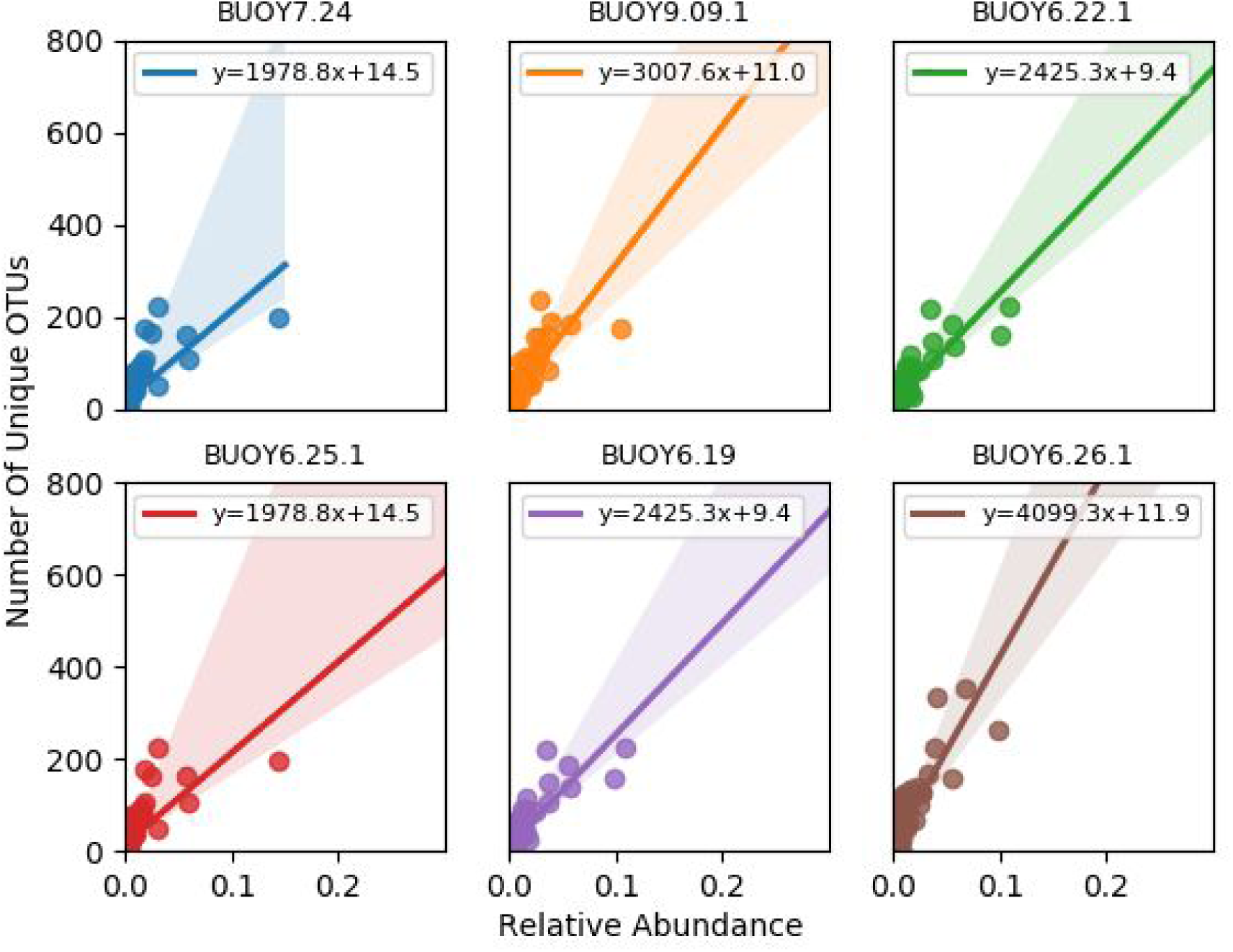
Diversity slopes per sample. Each sample from the Diversity Table is plotted with the number of unique OTUs vs. the relative abundance. Each sample has different slopes which can be used for further analysis.

After we are able to conduct a linear regression on the diversity table for each sample, we then plot the distribution of the slopes for all the samples in the BUOY site, shown in Figure 5. We can see from the histogram, which it is a skewed distribution with varying slopes of diversity.

**Figure 5.**
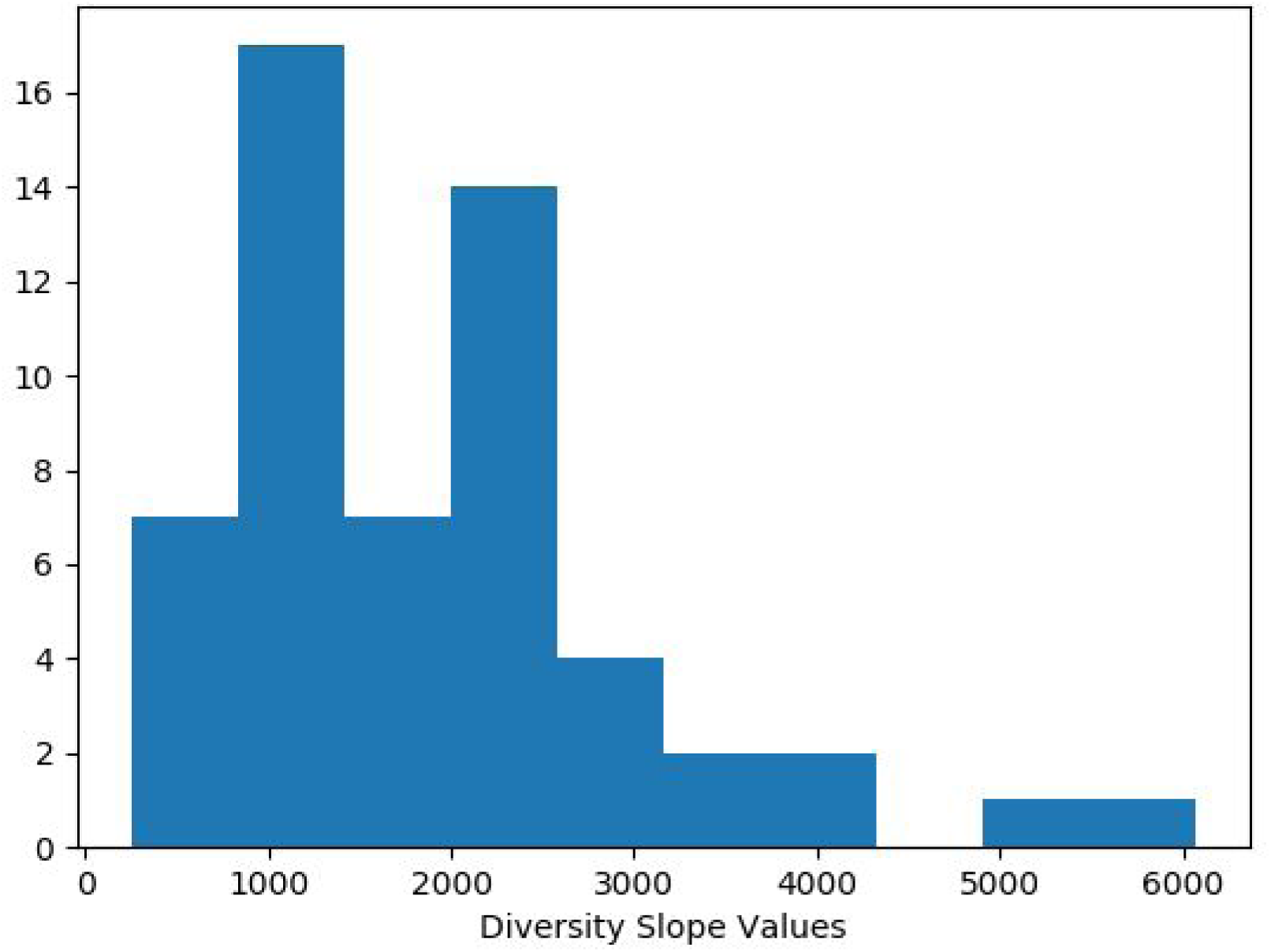
Histogram of Diversity Slope values. For all 55 samples from the buoy dataset, we plot the distribution of the diversity slopes.

We also conducted statistical analysis on the diversity slope values based upon the bloom stage. There are three bloom stages, Pre-Bloom, Bloom, and Post-Bloom. We group all the samples based on their bloom stages, and associate the sample diversity slope values. In Figure 6, we plot a boxplot for the diversity slope values for the three bloom stages. We can see that the distribution of slopes for each bloom stage is different.

**Figure 6.**
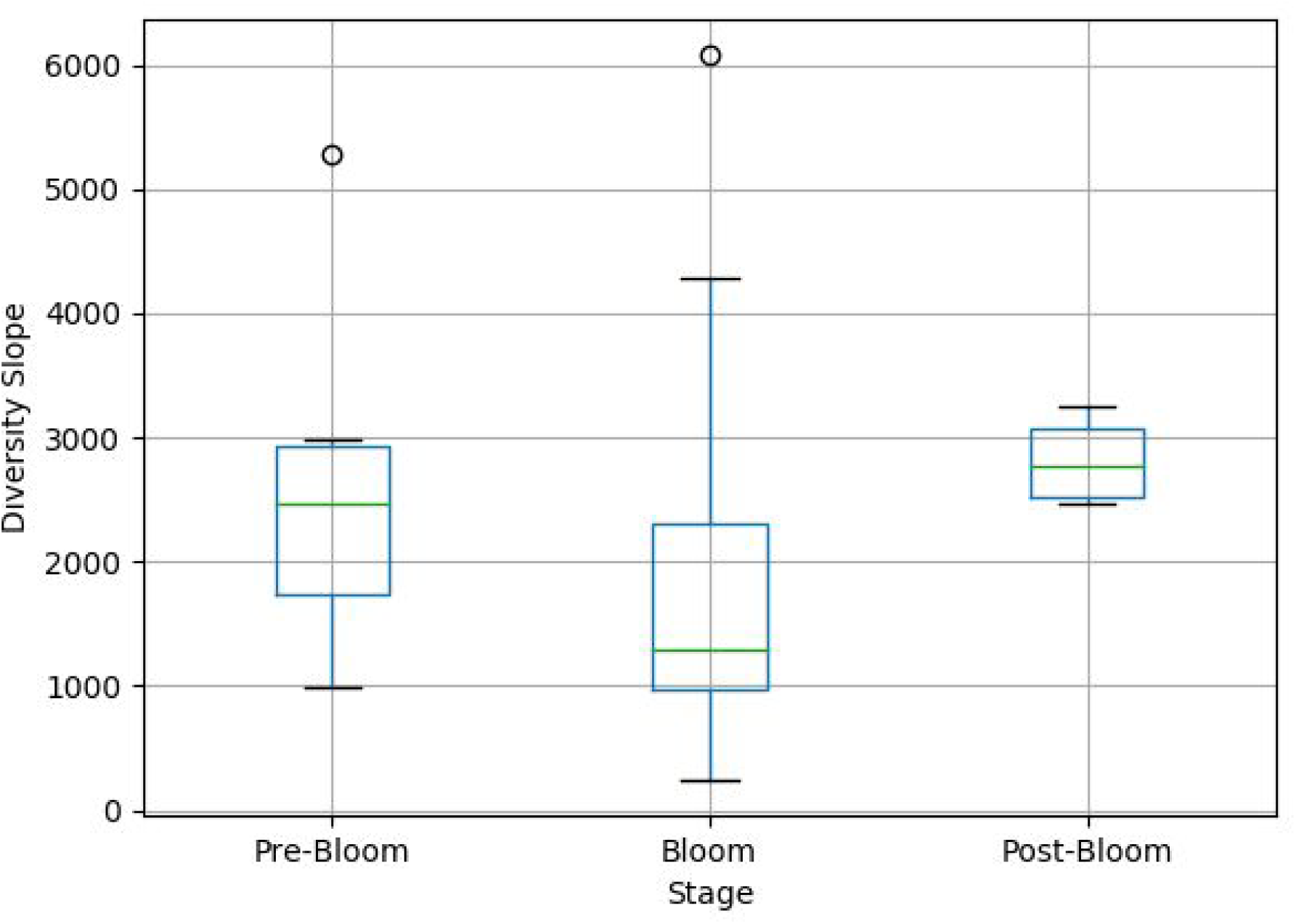
Boxplot of diversity slope values based upon bloom stage.

For further analysis, in Table 1, we conducted a Type-II ANOVA on the diversity slopes associated with their bloom stage. The p-value from the ANOVA gives 0.058906, which gives reasonable evidence that there is a difference in mean between the different bloom stages. This confirms the interpretation of varying distributions of slope values as seen in Figure 6. However, more analysis needs to be done, or more samples need to be analyzed as the number of samples used was 55, and one of the stages - ‘Post-Bloom’ - had 4 diversity slope values.

**Table 1.** Type-II ANOVA.

|  | sum_sq | df | F | PR( > F) |
| --- | --- | --- | --- | --- |
| C(Stage) | 8.110814e+06 | 2.0 | 2.991782 | 0.058906 |
| Residuals | 7.048682e+07 | 52.0 | N/A | N/A |

## Methods

Figure 1 describes our methodology in the very highest level. We start with an OTU table, which we then pick OTUs that higher than some threshold. At this step, our method is able to construct a new OTU table that combines redundant and trimmed OTUs. Then, this new OTU table is then used to construct a diversity table in which we focus on individual taxa and how the number of unique OTUs relate to the relative abundance of the taxa in a given sample.

In Figure 2, describes our methodology in a lower level. We start with the original OTU table. We then do an OTU seq retrieval from a sequence file that contains all the OTU sequences. We group the individual OTUs within their own associated Taxa. With each OTU grouped with their associated Taxa, we then use mafft for the multiple sequence alignment.

Once the OTU sequences are aligned, we then trim the sequences in the multiple sequence alignment such that the output becomes a group of subsequences that have high confidence (i.e sequences with no flanking gaps). The result of this can be exemplified in Figure 3. This is done first by converting the multiple sequence alignment into a matrix, which is filled with binary bits: 0s and 1s, where each 1 represents a gap, and 0 for others. The algorithm searches a trimming point from both sides to get rid of gaps that flank the ends of the alignment. One loop starts in the left end and then moves to the right until there are no gaps; the right side is processed similarly; the indices where there are no more gaps throughout the alignment are used to trim the alignments. This then creates a sub-alignment where another set of loops are made to ensure that at both ends there are two adjacent columns that have identical reads within the alignment.

We then filter out taxa where there is only one unique combined OTU.

Afterwards, we assign new taxonomy of the OTUs in the alignments by blasting these sequences against the Silva 16S rRNA gene database (Quast et al. 2013). Only the best hits for the OTU sequences are kept. When multiple best hits are returned, we keep them all if they have the same identity, alignment length, and E-value. Our tool provides a parameter called ‘similarity’ to specify a desired threshold as a cutoff to filter the hits to the Silva database that are below that threshold. The threshold used in our analysis is 97%.

We then create a new OTU table which contains the trimmed nonredundant OTUs with their combined relative abundance from the original OTU table and their new taxonomy. The OTUs with only one OTU for a taxa were filtered out and we add them to the new OTU table with ‘N/A’ to their new taxonomy as being not polymorphic, along with their sequence.

Lastly, from this new OTU table, we can construct another table that focuses on a specific sample. WITOD allows the ability to pick specific columns from the original OTU table, and complete the same procedure but in the final step be able to count the number of unique OTUs and associate it with the OTUs taxa. Also WITOD creates the cumulative relative abundance for the Taxa in the sample by adding up all the relative abundance for each of the unique OTUs in that Taxa. Thus the new table, called the Diversity Table for a sample has the information for what Taxa are in the sample, the cumulative relative abundance of that Taxa in the sample, and the number of unique OTUs for that specific Taxa.

